# Spatiotemporal Dynamics of Sound Representations reveal a Hierarchical Progression of Category Selectivity

**DOI:** 10.1101/2020.06.12.149120

**Authors:** Matthew X. Lowe, Yalda Mohsenzadeh, Benjamin Lahner, Ian Charest, Aude Oliva, Santani Teng

## Abstract

As the human brain transforms incoming sounds, it remains unclear whether semantic meaning is assigned via distributed, domain-general architectures or specialized hierarchical streams. Here we show that the spatiotemporal progression from acoustic to semantically dominated representations is consistent with a hierarchical processing scheme. Combining magnetoencephalography (MEG) and functional magnetic resonance imaging (fMRI) patterns, we found superior temporal responses beginning ~80 ms post-stimulus onset, spreading to extratemporal cortices by ~130 ms. Early acoustically-dominated representations trended systematically toward semantic category dominance over time (after ~200 ms) and space (beyond primary cortex). Semantic category representation was spatially specific: vocalizations were preferentially distinguished in temporal and frontal voice-selective regions and the fusiform face area; scene and object sounds were distinguished in parahippocampal and medial place areas. Our results are consistent with an extended auditory processing hierarchy in which acoustic representations give rise to multiple streams specialized by category, including areas typically considered visual cortex.

## INTRODUCTION

Hearing feels immediate and compulsory. In just a few hundred milliseconds, the human brain localizes sound sources (Wiegrebe, 2001), separates them from background noise and reverberation (Kell and McDermott, 2019; Teng et al., 2017; Traer and McDermott, 2016), determines their pitch (Norman-Haignere et al., 2019; Pressnitzer et al., 2001) and identifies them (Charest et al., 2009; Murray et al., 2006). While early acoustic attributes are invariably transformed into meaningful auditory object representations, the neural architecture and functional organization that facilitates auditory perception in humans remain unclear.

Unequivocal evidence of a general hierarchical functional organization in human audition has proved elusive. Despite considerable support for multistage hierarchical organization in nonhuman species (Kaas and Hackett, 2000; Kaas et al., 1999; Rauschecker and Tian, 2000; Rauschecker et al., 1995), the underlying functional architecture of the auditory system is less well established compared to that of the visual system, especially in humans, uniquely specialized for speech and music (McDermott, 2018; Norman-Haignere et al., 2019). Anatomically, primary auditory and early visual cortex are not directly analogous: peripheral input to PAC traverses more synapses compared to V1 (King and Nelken, 2009); and both PAC and nonprimary auditory areas, at least in monkeys and cats, receive direct medial geniculate projections (Hackett, 2011; Kaas and Hackett, 2000). Functionally, PAC BOLD responses resemble intermediate more than earlier layers of a task-optimized neural network (Kell et al., 2018), and A1 neuronal responses lack an obvious organizational axis beyond tonotopy (King and Nelken, 2009). Spectral and modulatory selectivity has been reported all along the STG, not just primary cortex, in both humans and nonhumans (Bizley and Cohen, 2013; King and Nelken, 2009; Staeren et al., 2009). The strongest and most abundant evidence for auditory cortical hierarchy comes from studies of speech processing (Evans and Davis, 2015; de Heer et al., 2017; Okada et al., 2010; Overath et al., 2015; Rauschecker and Scott, 2009; Yi et al., 2019), but this distribution does not necessarily imply sequential processing, nor does it necessarily generalize to other complex naturalistic auditory coding (Grady et al., 1997; Kell et al., 2018; Scheich et al., 1998; Sweet et al., 2005; Zatorre et al., 2002). Further, the anatomical, hodological, and neurophysiological data underpinning the primate core-belt-parabelt division in the auditory cortex (Hackett et al., 2001; Kaas and Hackett, 2000; Kaas et al., 1999; Rauschecker and Scott, 2009; Rauschecker and Tian, 2000; Rauschecker et al., 1995) are only sparsely available or inconsistent in humans (Hackett, 2011; McDermott, 2018; but see Nourski, 2017; Yi et al., 2019).

Hierarchies are spatial, temporal, and informational in nature: the sensory processing cascade propagates sequentially in parallel streams over multiple brain areas and hundreds of milliseconds, undergoing numerous representational transformations along the way (Bizley and Cohen, 2013; Rauschecker and Scott, 2009). Thus, characterizing hierarchy requires not only identifying multiple levels of neural representations over space, but also establishing their succession over time.

Here, we address this challenge via similarity-based fusion of human magnetoencephalography (MEG) and functional magnetic resonance imaging (fMRI) responses (Cichy and Oliva, 2020; Cichy et al., 2014, 2016a; Henriksson et al., 2019; Khaligh-Razavi et al., 2018; Mohsenzadeh et al., 2019; Salmela et al., 2018). Combining the spatiotemporal resolution of these imaging modalities within the representational similarity analysis (RSA) framework (Kriegeskorte and Kievit, 2013; Kriegeskorte et al., 2008), we hypothesized that (a) acoustic-dominated responses give way to category-dominated responses over both space and time, and (b) auditory semantic categories are selectively coded by high-level regions, rather than being distributed. To test these hypotheses, we analyzed cortical responses to naturalistic sounds of human voices, animal vocalizations, objects, and spaces at multiple representational levels, operationalizing their progression by comparing acoustic vs. semantic category-dominant coding. We found that semantic dominance increased systematically over time; that its spatial progression over regions of interest (ROIs) correlated with those ROIs’ peak response latencies; and that distinct high-level frontal, temporal, medial and occipital regions rapidly coded specific semantic categories. Our results support a hierarchical account of auditory object representations comprising parallel streams arising in core auditory cortex and rapidly targeting high-level auditory and multisensory regions.

## RESULTS

Human participants (n = 16) listened to 80 different monaural real-world sounds, presented diotically in random order for 500 ms every three seconds, while MEG and fMRI data were acquired in separate sessions. Stimuli were drawn from a large collection comprising sounds from animate (human and animal vocalizations) and inanimate (objects and scenes) sources. To dissociate acoustic from semantic category attributes, we equated root-mean-square intensities and spectral properties across categories (see **Methods** for details). Prior to the fMRI and MEG neuroimaging sessions, participants listened to each sound accompanied by a written description (see **Table S1**), so that the interpretation of each individual sound was not ambiguous. No explicit category information was provided. Participants were instructed to focus attentively on each sound for the duration of the trial and responded via button press to oddball sounds (200 ms pure tones presented in pairs, separated by 100 ms silence) presented an average of every ten trials. All participants performed at or near-ceiling on this vigilance oddball sounds task (mean hit rate 98.44% ± 1.38%, d’ = 5.71 ± .57).

### MEG-fMRI Fusion Reveals a Spatiotemporally Ordered Cascade of Neural Activation

To generate a spatiotemporally unbiased view of neural responses to sounds, we applied wholebrain searchlight-based fMRI-MEG fusion (Cichy et al., 2014, 2016a; Khaligh-Razavi et al., 2018; Mohsenzadeh et al., 2019), relating temporal neurodynamics in MEG with spatial BOLD activity patterns in fMRI. In brief, MEG and fMRI brain responses were decoded separately, then correlated within the RSA framework (Kriegeskorte et al., 2008). To this end, we first applied multivariate pattern analysis (MVPA) to MEG data in epochs spanning −200 to +3000 ms relative to stimulus onset. At each time point in the epoch, stimulus conditions were decoded pairwise, with classification accuracy indexing dissimilarity, to construct representational dissimilarity matrices (RDMs) (Cichy et al., 2014, 2016a; Kriegeskorte and Kievit, 2013). This yielded 3201 MEG RDMs of size 80 x 80 (1 per millisecond in the epoch). For fMRI, we used a whole-brain searchlight approach (Haynes and Rees, 2005; Kriegeskorte et al., 2006) and computed RDMs for each voxel (Kriegeskorte et al., 2008), using 1 – Pearson correlation as the pairwise dissimilarity metric, yielding a total of 40122 fMRI RDMs, also of size 80 x 80. Finally, we cross-correlated every MEG and fMRI RDM (Spearman ρ), then thresholded and cluster-corrected the resulting correlations (cluster-definition threshold *p* < 0.001, cluster size threshold *p* < 0.05; see Figure 1a and **Methods** for details). This yielded a series of whole-brain maps of MEG-fMRI correspondences across time, each serving as a frame in a “movie” spanning the epoch (see **Movie S1**).

**Figure 1.**
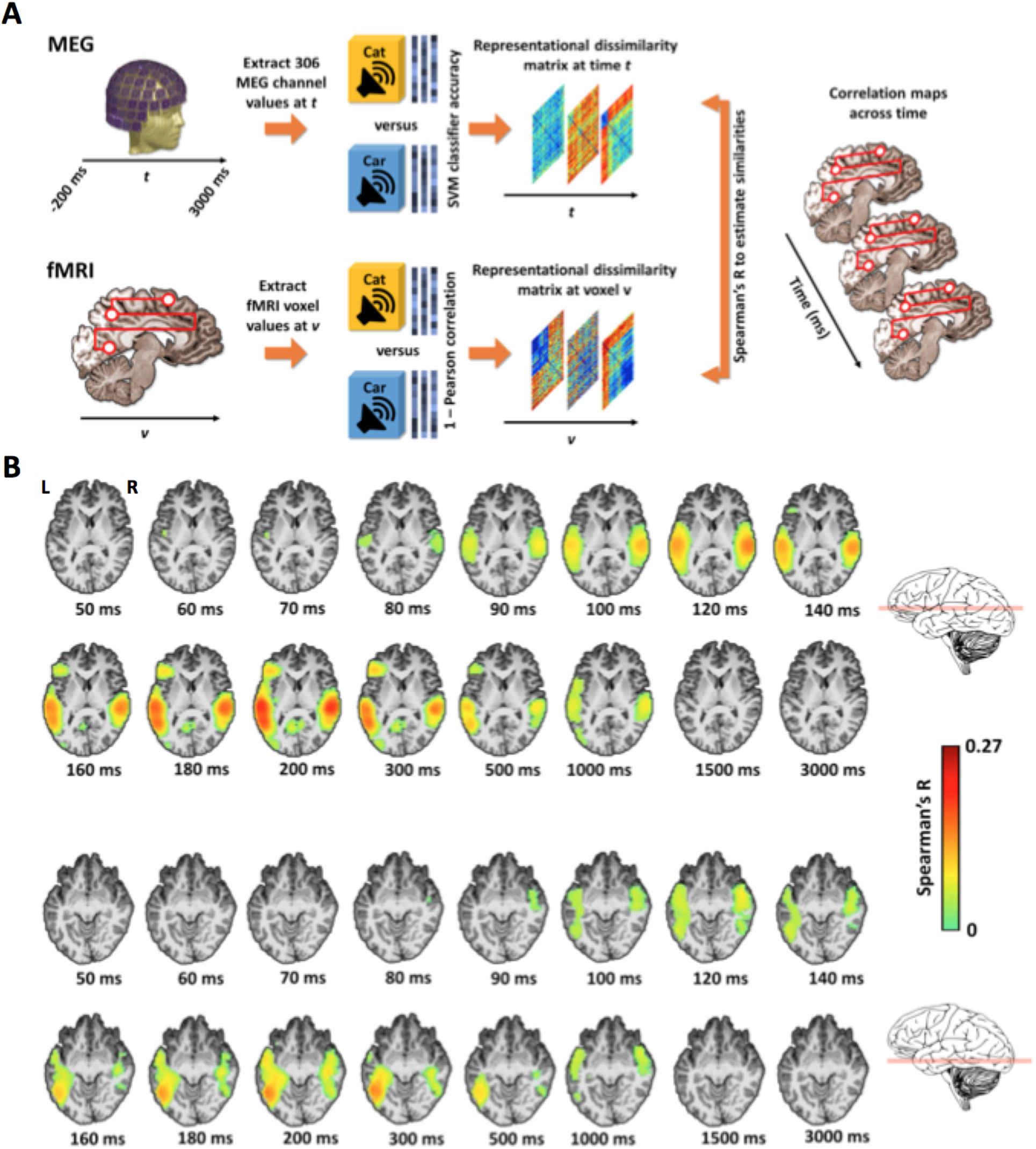
Spatiotemporal propagation of auditory neural responses indexed by MEG-fMRI fusion. **A.** Overview of MEG-fMRI searchlight approach. Brain responses acquired in separate sessions for each subject were analyzed to generate representational dissimilarity matrices (RDMs) over time (MEG) and space (fMRI), then cross-correlated to produce whole-brain correlation maps across time. **B. Main cortical regions involved in the searchlight-based fMRI-MEG fusion.** Illustration on two axial slices (z = 10 and −10 mm) at select time points. There are significant neural representations beginning over the auditory cortex and spreading towards pre-frontal, ventral and medial regions. Color-coded voxels indicate the strength of MEG-fMRI RDMs correlations (Spearman’s *R*, scaled between 0 and maximal observed value, n = 16, cluster definition threshold p < 0.001, cluster threshold p < 0.05). See the fullbrain fusion in **Movie S1.**

Significant fusion correlations appear along the superior temporal gyrus starting 80-90 ms post-stimulus onset, spreading anteriorly and posteriorly along the superior temporal plane, and reaching pre-frontal, ventral occipitotemporal, and medial regions by ~130-140 ms (see **Figure 1b**, axial slices at z = 10 and −10 mm). Whole-brain fusion thus illustrates an orderly spatiotemporal progression of responses from early sensory cortices to higher-level and extratemporal regions, consistent with processing streams in hierarchical organization (Rauschecker and Scott, 2009).

We further quantified the spatiotemporal distribution of the brain response by repeating the fusion approach in a region-of-interest (ROI) analysis, which yields a separate fusion correlation time course for specific cortical regions. While we hypothesized a progression in which MEG-fMRI correspondence appears earliest in core auditory ROIs and travels systematically outward, the nature and organization of this progression remains unclear. We defined primary and nonprimary auditory anatomical ROIs spanning the superior temporal gyrus (STG) and inferior frontal gyrus (IFG), and functionally defined occipital and temporal ROIs known to be category-selective for faces, objects and scenes in vision as well as audition.

Specifically, the ROIs comprised the primary auditory cortex (PAC, composed by TE1.0 and TE1.1), TE1.2, planum temporale (PT) and planum polare (PP) (Desikan et al., 2006; Morosan et al., 2001; Norman-Haignere et al., 2013). From Pernet et al. (2015), we identified the voice-selective left inferior frontal gyrus (LIFG) region, and a modified temporal voice area (designated TVAx, with voxels overlapping with PT removed). We also selected the Parahippocampal Place Area (PPA, Epstein and Kanwisher, 1998), the Medial Place Area (MPA, Silson et al., 2016), the Fusiform Face Area (FFA, Grill-Spector et al., 2004; Kanwisher et al., 1997), and the Lateral Occipital Complex (LOC, Grill-Spector et al., 1999; Malach et al., 1995). Finally, we defined the Early Visual Cortex (Lowe et al., 2016; EVC, MacEvoy and Epstein, 2011) as a control region (see **Methods** for details).

Similarly to the whole-brain searchlight analysis, for each participant and ROI, we extracted voxelwise fMRI BOLD responses and computed an RDM from pairwise Pearson correlation distances. We then correlated each group-averaged ROI RDM with subject-specific MEG RDMs over time to generate an fMRI-MEG fusion correlation time course per ROI per subject.

As shown in Fig. 2 and Table 1, statistically significant clusters (defined using sign-permutation tests, *p* < 0.01 cluster-definition threshold, *p* < 0.05 cluster threshold) were observed in all ROI fusion time courses (except EVC). Early primary and non-primary auditory ROIs (PAC, PT, TE1.2, PP) exhibited similar peak latencies of ~115 ms post-stimulus onset. Peaks in voice-selective regions (TVAx and LIFG) and MPA occurred at ~200 ms, and ~300 ms or later in the other ventral and medial functionally defined regions (FFA, PPA, and LOC). We further characterized the timing of the processing cascade by comparing the MEG–fMRI fusion time course peaks between PAC and the other ROIs (see Table 1, latency difference). Compared with PAC, MEG–fMRI correspondence in representations emerged significantly later (at least 80 ms) in the voice, face, object and scene-selective regions. These results confirm that sound-related neural activity in higher cortical regions occurred later than in the early auditory regions, corroborating and quantifying the signal propagation revealed in the whole-brain searchlight analysis.

**Table 1.** Mean first-peak latency for 10 ROIs and comparison of peak latencies of PAC versus ROIs for interval [-200, 1000] ms. All values are averages across subjects (n=16) with 95% confidence intervals in brackets. Latency differences from PAC and significance were determined by bootstrapping (5000 times) the sample of participants (1000 samples). Cluster size threshold *P*= 0.05, significance threshold *P* = 0.01

| Region of Interest | Peak latency (ms) | Latency difference (ms) | Significance (*P* value) |
| --- | --- | --- | --- |
| PAC (TE1.0 & TE1.1) | 115 (100, 191) | - | - |
| TE1.2 | 117 (111, 207) | 2 | n.s |
| Planum Temporale (PT) | 115 (103, 197) | 0 | n.s |
| Planum Polare (PP) | 117 (110, 207) | 2 | n.s |
| Temporal Voice Area (TVAx) | 199 (188, 251) | 84 | 0.006 |
| Left Inferior Frontal Gyrus (LIFG) | 196 (188, 380) | 81 | 0.003 |
| Fusiform Face Area (FFA) | 298 (200, 368) | 183 | 0.0006 |
| Parahippocampal Place Area (PPA) | 293 (219, 380) | 178 | 0.004 |
| Medial Place Area (MPA) | 199 (196, 400) | 84 | 0.001 |
| Lateral Occipital Complex (LOC) | 377 (169, 410) | 262 | 0.008 |

**Figure 2.**
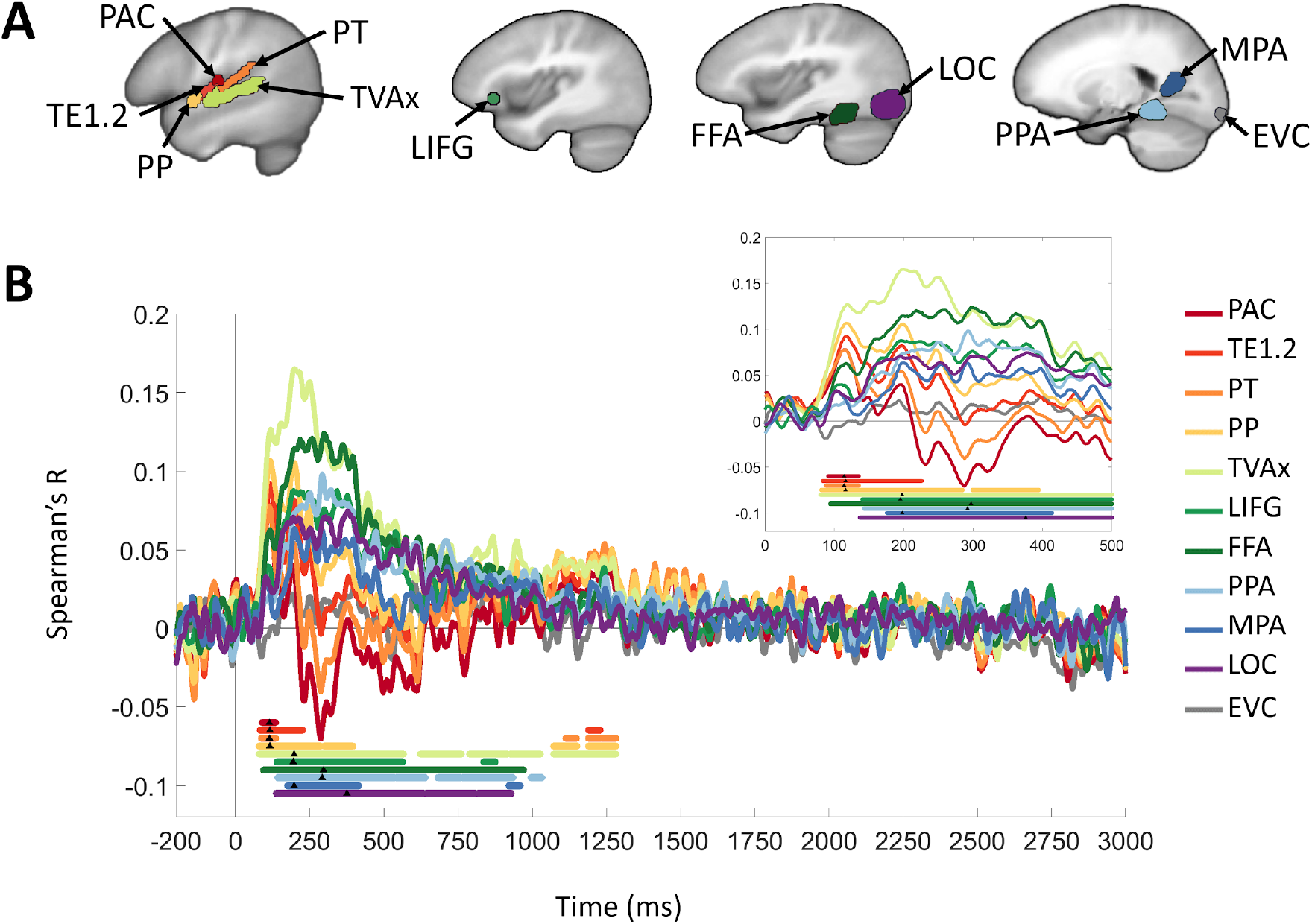
ROI-based MEG-fMRI fusion time courses. **A.** Eleven regions of interest (ROIs) spanning temporal, frontal, and occipital lobes were selected to examine auditory responses, including the primary auditory cortex (PAC), TE1.2, planum temporale (PT), planum polare (PP), temporal voice area (TVAx), left inferior frontal gyrus (LIFG), fusiform face area (FFA), parahippocampal place area (PPA), medial place area (MPA), lateral occipital area (LOC), and early visual cortex (EVC). **B**. ROI-specific fusion time courses (color-coded to match ROI illustrations in A) indexing correlation between whole-brain MEG and a single RDM representing each fMRI ROI. Gray vertical line denotes stimulus onset at t=0 ms. Solid color-coded bars below time courses represent significant time points, observed for all regions except EVC; black triangles indicate respective peak latencies. Inset magnifies 0-500 ms regime (the duration of the stimulus) for clarity. All statistics, P<0.01, C<0.05, 1000 permutations.

### Systematic Trend From Acoustic to Semantically Dominated, Category-Specific Coding

While the fusion analysis indexes the propagation of spatiotemporal neural response patterns, it is agnostic to their representational content. A hierarchical account of auditory processing would predict that such a cascade initially encodes acoustic, then progressively higher-level stimulus representations (Bizley and Cohen, 2013; Rauschecker and Scott, 2009).

To capture this transition, we operationalized representational levels at two extremes: First, we constructed a *Cochleagram* RDM comprising euclidean distances between stimulus log-frequency cochleagrams (see **Methods** and **Figs. 3 & S1**) to model the hypothesized low-level similarity structure of auditory afferents arriving from the cochlea. Second, we devised a *Category* RDM hypothesizing a generalized high-level semantic category selectivity across all four categories: all within-category pairwise distances were set to 0, and all between-category distances to 1. The strength of these representational levels was indexed by computing Spearman correlations between the models and brain data.

**Figure 3:**
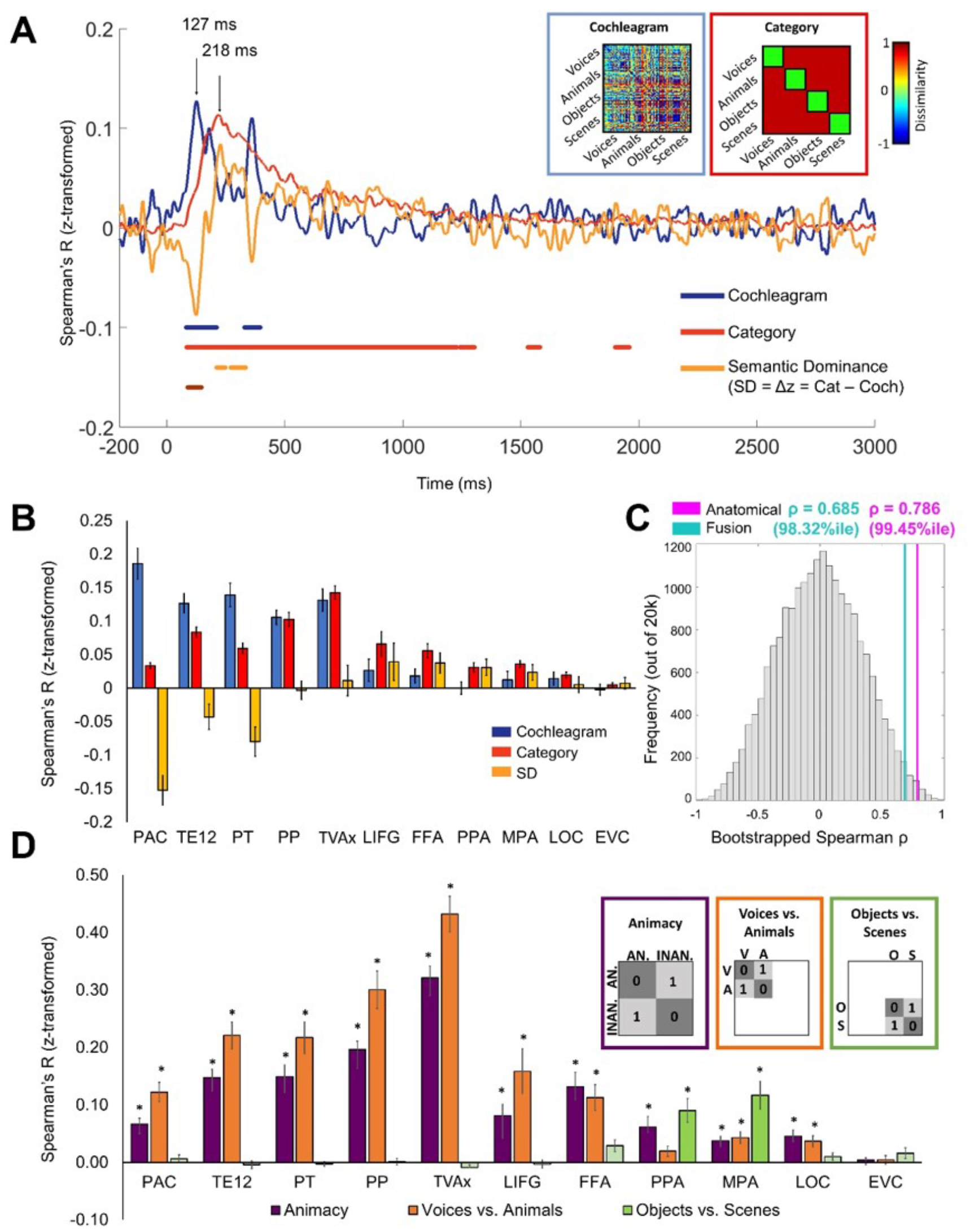
Emerging category vs. acoustic selectivity across time and ROIs. **A.** Temporal dynamics in MEG responses: Cochleagram (blue) and Category (red) model RDMs shown in insets encode hypothesized similarity structures of acoustic and semantic neural stimulus representations. Pairwise Cochleagram model dissimilarities are indexed by the euclidean distance between stimulus log-frequency spectra; for the Category model, all between-category pairs were assigned a value of 1 and within-category pairs a value of 0. Plot shows Fisher z-normalized correlation time courses between models and whole-brain MEG data. Cochleagram and Category peaks at 127 (CI 119–178) ms and 217 (CI 209–259) ms, respectively. Yellow trace is Semantic Dominance (SD) difference score, Δ*z* = *z*_Category_ *z*_Cochleagram_. Negative SD = acoustically dominated responses; positive SD = semantic category-dominated responses. Color-coded bars indicate significant temporal clusters for each trace. **B.** Model correlations with RDMs computed from fMRI ROI data. Color code same as in **A**. Bar plots depict the subject-averaged measure in each ROI; error bars = standard error of the mean (SEM). Bonferroni corrected one-tailed t-test, asterisks indicate significance at p <.05. **C.** Histogram of 20,000 bootstrapped ROI-SD trend rank correlations compared with empirical correlations from our anatomical (magenta; 99.5%ile) and fusion-derived (cyan; 98.3%ile) ROI rank assignments. **D.** Semantic subcategory distinctions in fMRI ROIs. Subject-averaged bar plots indicate distinction between category animacy (purple), human vs. animal vocalizations (orange), and objects vs. scenes (green). Plotting conventions and statistics same as in **B**.

While our stimulus set was designed to minimize confounds between these models (see **Methods**), we would not necessarily expect a given set of fMRI voxels or MEG sensor patterns to correlate only with one and not the other, both due to the intrinsic correlation between sound acoustics and categories (Theunissen and Elie, 2014) and the fact that neurons throughout primary and nonprimary auditory cortex are responsive to low- and higher-level sound properties (King and Nelken, 2009; Norman-Haignere and McDermott, 2018; Staeren et al., 2009) However, in a hierarchical processing stream, we would expect their *respective* contributions to vary systematically between regions over time. Thus, we computed the difference between Category and Cochleagram model correlations (Fisher-z normalized to enable direct comparison) with MEG and fMRI patterns. This measure, which we term *Semantic Dominance* (SD), quantifies the relative weight of neural representations: a negative value indicates predominance of acoustic properties, and a positive value indicates predominance of semantic properties. We first evaluated the Cochleagram and Category models (and SD) separately on temporal and spatial data, then in a combined fashion guided by the fusion analysis.

Cochleagram and Category models both correlated significantly with the whole-brain MEG signal (Fig. 3A). The correlation time courses reached significance at similar onset times of ~80 ms; the period of significant Cochleagram model correlation was shorter and had an earlier peak latency (127 ms; 95% confidence interval 119–178 ms) compared to the Category model, which peaked at 218 (209–259) ms and remained significant through ~1300 ms. Semantic Dominance was significantly negative from ~90–150 ms and significantly positive from ~200– 350 ms. These results indicate a rapid onset of frequency-sensitive coding, a broad temporal regime of general semantic category-sensitive coding, and a clear temporal progression in which neural response patterns are more differentiable by semantic category than by their spectra after about 200 ms.

The Cochleagram model correlated significantly with all superior temporal fMRI ROIs (PAC, TE1.2, PT, PP, TVAx), with the PAC correlation significantly higher than the nonprimary ROIs (all p < 0.05). Frontal (LIFG) and functionally defined (FFA, PPA, MPA, LOC, EVC) ROIs showed no significant Cochleagram correlation. By contrast, the Category RDM correlated significantly with all ROIs except EVC (Fig. 3B).

To quantify the relationship between these results, we next tested for a systematic trend in the relative strengths of representational levels by computing a Spearman rank correlation between ROIs and their Semantic Dominance score. Our reasoning, adapted from a similar approach in prior work (Fischer et al., 2011), was that the ROI ranks reflect our hypothesized ordering schemes, and that a significant positive correlation would indicate a systematic trend along the ranking dimension. We therefore assigned ranks based on independent spatial (functional anatomy) and temporal (fusion-derived peak latency) criteria. For the spatial analysis, we first differentiated primary auditory cortex based on convergent anatomical, histological and functional criteria (Morosan et al., 2001; Norman-Haignere and McDermott, 2018; Sweet et al., 2005), then ranked non-primary areas progressing along and beyond the supratemporal plane. We assigned PAC a rank of 1; TE1.2, PP, and PT (all adjacent to PAC) a rank of 2; TVAx (farther along the posterolateral STG) a rank of 3; LIFG a rank of 4; and FFA, PPA, MPA and LOC a rank of 5. (EVC was excluded, as no significant model correlations were found in that ROI.) The trend was significantly positive (ρ = 0.786, p = 0.0071), indicating that an independent anatomically based ranking closely tracks increasingly category-selective neural representations.

To test our temporally based ordering scheme, we repeated this analysis, this time ordering ROIs guided by their empirically determined fusion peak latencies (as shown in Table 1/Fig. 2b). Previous work (Cichy et al., 2016a) suggests that correlation peak, rather than onset, latencies more closely track electrophysiologically measured neuronal dynamics. As before, the correlation (ρ = 0.685, p = 0.0288) indicates a significant positive trend in Semantic Dominance across ROIs ranked by an independent empirical metric of their fusion correlation peaks; i.e., neural responses become dominated by category-selective patterns over time.

While our rankings were guided by these criteria, they may reflect hidden factors or correlate spuriously with the brain data. Thus, to test our *a priori* anatomical and empirical ROI-fusion-derived ranking schemes against all possible ROI rankings, we compared our ROI-rank-vs.-Semantic-Dominance correlations from those schemes against a 20,000-sample bootstrapped distribution of rank correlations, each computed by assigning each ROI a randomly drawn rank between 1 and 10 (with replacement). The anatomical and fusion-timing-based correlations were higher than 99.5% and 98.3% of all bootstrapped values, respectively (Fig. 3C), indicating that our independently computed Semantic Dominance measure closely matches activation patterns in ROIs ranked by anatomical outward progression from PAC, as well as by the temporal sequence of peak fMRI-MEG correspondences in those ROIs.

The SD trend shows that pooled across multiple categories, semantic relative to acoustic selectivity increases over time and space. However, it could conceivably index a distributed, generic representation of high-level stimulus attributes, whereas modular organization into disparate maps and streams (Rauschecker and Scott, 2009) would predict differential category selectivity across ROIs. To disambiguate between these possibilities, we examined the specificity of semantic coding by partitioning the fMRI RDMs into pairwise between- and within-category portions, with the average differences between correlation distances indexing category distinctions. In this way we parcellated the generic Category model to investigate broad animacy selectivity (Voices & Animals vs. Objects & Scenes) as well as more granular sensitivity to differences between animate (Voices vs. Animals) and inanimate (Objects vs. Scenes) categories. As shown in Fig. 3D, Human and animal vocalizations were best differentiated (more than general animacy) in superior temporal and voice-selective ROIs, while inanimate environmental sounds were chiefly differentiated in scene-selective PPA and MPA, suggesting functional specialization of auditory responses in these areas.

## DISCUSSION

Here we analyzed the spatial, temporal, and representational structure of cortical responses to naturalistic sounds in human listeners. We leveraged the temporal resolution of MEG and the spatial resolution of fMRI, fusing them within the RSA framework to track the processing cascade from early auditory cortex to frontal, ventral, and medial occipital regions. The whole-brain searchlight analysis revealed an orderly progression of neural activation originating in bilateral temporal lobes and spreading anteriorly and posteriorly to extratemporal regions throughout much of the cortical volume. The timing of MEG-fMRI correspondence in specific regions of interest bears this out concretely, with the earliest peak latencies in superior temporal ROIs (PAC, TE1.2, PT, PP), followed by voice-(LIFG, TVAx) and scene-selective (MPA) regions and reaching ventro-temporal and occipital category-selective regions (FFA, PPA, LOC), but not early visual cortex. The representational content of the response patterns was initially strongly biased toward low-level acoustic features, as indicated by strong correlations with the Cochleagram model of auditory nerve afferents, but increasingly dominated by high-level semantic categories, as indicated by correlations with the Category model RDM. The Semantic Dominance difference score indexed the systematic trend between these two representational extremes over time (significant sign reversal of SD over the MEG epoch) and space (significant positive trend of SD across fMRI ROIs). The ROI rankings modeling the SD trend were independently estimated from two metrics: *a priori* functional-anatomical layout and empirically determined MEG-fMRI fusion time courses. These rankings produced higher trend correlations than almost all possible rankings, supporting the robustness of our convergent temporal and spatial bases for quantifying the evolving nature of the processing stream. Finally, more granular interrogation of category selectivity revealed its spatial specificity, arguing against a generalized, distributed model of categorical sound representation. Taken together, our findings provide converging evidence for large-scale hierarchical organization of the human auditory stream through distinct levels of processing dissociable by space, time, and content.

### An Integrative Approach to Identifying Attributes of Hierarchical Coding

A hierarchical processing model posits pathways that, in parallel, transform “low-level” sensory input serially into increasingly complex, abstract “high-level” representations. Consequently, it is reasonable to expect that the physical substrate of these pathways would have sequential properties as well: as the temporal dynamics of representations follow a low-to-high-level sequence, increasing distance from a low-level origin should track with higher-level representations, which are routed to distinct loci. We based our hypotheses on these principles, operationalized for the auditory system (Bizley and Cohen, 2013; Kell et al., 2018; Rauschecker and Scott, 2009).

Our integrative approach addresses several methodological and conceptual barriers to resolving conflicting interpretations arising from the body of earlier work. First, the temporal specificity that MEG-fMRI fusion adds to spatially mapped brain responses allows us to directly test spatial hypotheses to which time is intrinsic, such as sequential processing streams carrying evolving levels of representation. In this way, the fusion analysis provides neural data at a level typically unavailable in humans. Even the human intracranial literature, while largely bearing out our results (Nourski, 2017), investigates higher-level processing narrowly with speech (Sahin et al., 2009; Steinschneider et al., 2014), and does not provide the full-brain coverage of fMRI/MEG. Second, beyond enabling the fusion analysis, the RSA framework allows us to operationalize high- and low-level representations of a large naturalistic stimulus set spanning a range of commonly encountered sound classes. Finally, the Semantic Dominance difference score analysis allows us to manage the confounds between acoustic and semantic properties of naturalistic sounds (Norman-Haignere and McDermott, 2018), and to test explicitly for spatial and/or temporal trends in their relative contributions to neural response patterns.

### Separating Levels of Representation

How should we interpret the transition from Cochleagram to Category model dominance? Our modeling analyses capture two endpoints of a neural processing cascade transforming a waveform to a semantically organized representation. Other models representing hypothesized processing stages could be operationalized, e.g., as neuronal responses to different acoustic properties (Rauschecker and Scott, 2009) components of complex sounds (Teng et al., 2017), anatomical location (Norman-Haignere and McDermott, 2018), or layers of task-optimized neural networks (Kell et al., 2018). The RSA framework and our analysis are adaptable to any of these approaches. Indeed, very recent work mapped multivariate MEG responses to various spectrotemporal properties of natural sounds (Ogg et al., 2020), but could not specify the extent to which those decoding results show the temporal emergence of object-level semantic labels vs. the evolving grouping of acoustic response properties, as the stimulus set’s semantic and acoustic properties were not independent. As with visual images (Oliva and Torralba, 2001), sounds in the natural world contain some categorical structure in their low-level features (Charest et al., 2009; Theunissen and Elie, 2014), which complicates the separation of acoustic-vs. category-based neural decoding. We guarded against this confound by preparing the stimulus set to contain minimal category structure (See Methods and Fig. S1). Thus, the Category RDM was less likely to spuriously capture acoustic differences, but was additionally normalized relative to the Cochleagram RDM via the Semantic Dominance difference score. This approach allowed us to examine the same MEG or fMRI data for low- and high-level selectivity, and to quantify which selectivity dominates coding in a given area (and how that dominance changes). For example, PT and LIFG were each significantly sensitive to category and acoustic structure (Fig. 3B), but to opposite relative extents.

The widespread category selectivity we found may be reminiscent of modulation selectivity, likewise considered a widespread response property to model auditory cortical responses (Chi et al., 2005; Santoro et al., 2017). Could our results be spuriously indexing the emergence of modulation sensitivity? While it is impossible to rule out every possible correlate of a hypothesized pure stimulus semantic label, we consider it unlikely. First, the ubiquity of modulation selectivity in the auditory cortex may be overstated. The widespread fMRI responses elicited beyond PAC by natural sounds are not replicated by synthetic modulation-matched sounds (Norman-Haignere and McDermott, 2018), suggesting that nonprimary neurons may be more selective for other properties that co-occur with modulation. Second, that falloff in non-PAC modulation sensitivity contrasts with the increase in category sensitivity (especially relative to frequency coding) that we found both over time and with greater separation from PAC.

### Hierarchy Within and Beyond Superior Temporal Regions

A putative core-belt-parabelt tripartite hierarchy in human audition remains the subject of ongoing debate. We found functional differences between primary and early nonprimary ROIs along the STG: spectral sensitivity was significantly reduced and category sensitivity increased outside PAC (Figs. 3B,D). Our fusion time course results did not reveal significant latency differences among these ROIs (Fig. 2, Table 1). The overlapping onset and peak timing between PAC, TE1.2, PT, and PP may reflect parallel processing, possibly mediated by direct medial geniculate projections not only to core, but also belt and parabelt areas (Hackett, 2011; Kaas and Hackett, 2000). However, our data also reflect the aggregate responses to a large set of diverse, complex sounds. Our stimulus set and analyses controlled duration, mean intensity, and overall spectral structure across exemplars and semantic categories, but these still vary dynamically over the course of the sound and are known to modulate early cortical latencies in both neurophysiological and M/EEG studies (Biermann and Heil, 2000; Lakatos et al., 2005; Murray et al., 2006). Thus, a hierarchical architecture encountering many naturalistic sounds, with a variety of attack intensities, dominant frequencies, and bandwidths, would be expected to process them at variable speeds. Aggregated across the stimulus set, this could predict the overlapping early latencies we found, before the processing streams become more segregated in the wider cortical network.

Beyond superior temporal regions, fusion and model analyses show responses further separated by time, space, as well as content. The spatiotemporal progression of fusion responses tended toward progressively higher onset/peak latency and duration farther from PAC (Figs. 1, 2; Table 1), in accordance with the “multiple maps and streams” feature characteristic of a distributed hierarchy (Rauschecker and Scott, 2009). Superior temporal and extratemporal ROIs (LIFG, FFA) showed a clear sensitivity to stimulus animacy and, specifically, human vs. animal vocalizations (Fig. 3D, S2); fusion correlations for LIFG and TVAx both peaked at ~200 ms, a nearly exact match to frontotemporal voice-selective ERP peaks (Charest et al., 2009) and consistent with the notion of a functionally specialized rapid voice- and speech-processing network (Belin et al., 2000; de Heer et al., 2017; Norman-Haignere et al., 2019; Pernet et al., 2015). Notably, response patterns in FFA, typically considered a visual face-selective region, distinguished animate vs. inanimate and human vs. animal sounds comparably to the classical voice-processing network, with a later fusion peak but onset comparable to the superior temporal ROIs (Fig. 2B). Voices unconnected to specific identities typically elicit no or weak FFA activity at best (e.g. de Heer et al., 2017, in which FFA voxels responded to semantic but not spectral or articulatory aspects of speech); however, face and voice regions share functional and structural connectivity (Blank et al., 2011; Kriegstein et al., 2005), and nonvisual face-related stimuli elicit FFA-colocalized responses in congenitally blind persons (van den Hurk et al., 2017; Murty et al., 2020). Thus, the FFA selectivity (and rapid response compared to its anatomical neighbors) in our results may reflect its role in a person/identity-recognition mechanism with unified coding principles (Yovel and Belin, 2013) rather than voice processing *per se*.

Interestingly, only extratemporal regions, typically considered part of the visual ventral stream, distinguished between inanimate (scene and object) categories of sounds, with representations consistently correlating with semantic properties but only weakly or not at all to acoustic properties (Fig. 3B,D). Traditionally sensory-specific areas have long been implicated in cross-sensory processing (Jung et al., 2018; Kim and Zatorre, 2011; Kriegstein et al., 2005; Smith and Goodale, 2015; Vetter et al., 2014); however, in this case, the regions were activated in the absence of any visual stimulation or association with a visual cue, ruling out a multisensory integrative or paired associative process. Auditory objects and scenes have resisted easy analogical transfer from their visual counterparts, both as fundamental concepts (Griffiths and Warren, 2004) and as variables in experiments, due to differences in spatiotemporal structure of an image vs. a sound (Teng et al., 2017).

Consequently, could these category-specific representations in visually selective regions simply result from visual mental imagery? Substantial work documents the involvement of early visual cortex in mental imagery (Kosslyn et al., 1995, 2003); participants instructed to imagine sound scene categories without visual stimulation generate category-specific EVC representations (Vetter et al., 2014). Yet we found no evidence for category selectivity specifically, or MEG-fMRI correspondence generally, at any time points in EVC (Figs. 2,3, Movie S1). Still, imagery may occur without EVC activation, generating higher-order representations (Reddy et al., 2010). These representations emerge much later compared to feedforward signals. Cued visual imagery of faces and houses becomes decodable in brain dynamics after ~400 ms, peaking at ~1000 ms (Dijkstra et al., 2018); even brief, highly overlearned stimuli like auditory letter phonemes elicit visual imagery responses peaking at ~400 ms or later (Raij, 1999; Raij et al., 2000). By contrast, MEG-fMRI fusion latencies in most of the ROIs in our study peaked before 300 ms, with LOC peak latency at 377 ms; the outermost bounds of the 95% confidence intervals were 410 ms for LOC and 400 ms for MPA (Fig. 2B, Table 1). In fact, MPA peaked relatively early, at ~200 ms, a time scale for rapid auditory scene-object distinction on par with that of voice processing. The stimuli themselves were 500 ms in duration; a semantic understanding and resultant unprompted crossmodal imagery would be unlikely to arise and peak while the sound was still being presented to the listener.

Our overall results are therefore consistent with the accumulating evidence for a functional primary/nonprimary distinction in humans, inconsistent with an imagery account of higher-level representations, and extend previous work by tracing out an acoustic-to-semantic processing hierarchy from primary auditory, nonprimary auditory, and multiple high-level extratemporal cortical regions.

### Limitations and Future Directions

RSA-based M/EEG-fMRI fusion is constrained by some fundamental and practical limitations (cf. Cichy and Oliva, 2020). For example, its central assumption of isomorphism between neuromagnetic and BOLD responses is most likely to reveal aspects of neural signals accessible to both imaging modalities, and to raise the combined signal-to-noise threshold relative to each modality alone. In our study, we mitigated these limitations by testing our hypotheses via a combination of unimodal and multimodal analyses (Figs. 2,3). Additionally, the correlation of whole-brain MEG data with a single static map of fMRI data means that, theoretically, the dynamics of similar representations in two different regions cannot be distinguished. However, this issue can be overcome with carefully designed experimental paradigms and stimulus sets: our present study specifically maximized differences in representations of the stimulus set, both between low- and high-level patterns as well as different category-specific patterns.

These constraints notwithstanding, the fusion approach, primarily applied to visual systems to date, has proven a powerful way to elucidate spatiotemporal features of visual processing such as V1-Inferior-Temporal (IT) timing and evolution (Cichy et al., 2014, 2016a), scene layout encoding timing in occipital place area (OPA) (Henriksson et al., 2019), ventral-dorsal dynamics (Cichy et al., 2016a), task and attentional modulations (Hebart et al., 2018; Salmela et al., 2018), dissociable object size and animacy selectivity (Khaligh-Razavi et al., 2018), model-or behaviorally based similarity (Cichy et al., 2016b, 2019), and feedforward-feedback interplay (Mohsenzadeh et al., 2018) when viewing visual objects. The versatility of its framework is readily generalizable to outstanding questions in auditory neuroscience (cf. Cichy and Teng, 2017). For example, in the present study, we interrogated the generalized structure governing the neural representation of auditory category information; future variations on that paradigm could examine, e.g., task-contingent modulations of timing and stimulus representation in responses, representations drawn from behavioral tasks, or complex manipulations within a single domain such as speech, in which spatiotemporally resolved data has previously been available only via rare, invasive clinical procedures (Martin et al., 2018; Nourski, 2017; Sahin et al., 2009; Yi et al., 2019). Similarly, in addressing the ongoing debate over proposed dual “what” and “where” auditory pathways (Bizley and Cohen, 2013; Lomber and Malhotra, 2008; Rauschecker and Tian, 2000), multilevel response patterns to a large stimulus set that includes spatial as well as semantic manipulations could identify distinct and overlapping components of the processing streams from complex real-world stimuli. Further, the attentional, grouping, and segregation processes mediating auditory scene analysis (Lakatos et al., 2013; Shamma et al., 2011) could be parcellated with fine-grained spatial and temporal resolution. In all these cases, RSA-based M/EEG-fMRI fusion provides a powerful, conceptually straightforward integrative framework for analyzing the neural correlates of auditory perception and cognition.

## METHODS

### Participants

Sixteen right-handed, healthy volunteers with normal or corrected-to-normal vision and no hearing impairments (8 male, age: mean ± s.d. = 28.25 ± 5.95 years) participated in the experiment. The study was conducted in accordance with the Declaration of Helsinki and approved by the MIT Committee on the Use of Human Experimental Subjects. All participants completed one MRI and one MEG session. All participants provided written consent for each of the sessions.

### Stimuli

An initial set of 200 naturalistic monaural sounds was resampled to 44.1 kHz, root-mean-square normalized, and trimmed to 500 ms duration, including 10 ms linear rise and fall times. We computed cochleagrams for each sound using a Matlab-based toolbox (Brown and Cooke, 1994) that emulates the filtering properties of the human cochlea by transforming time-domain signals into a time-frequency representation of the average firing rate of auditory nerve fibers. Specifically, each waveform was divided into 1-ms bins and passed through a gammatone filterbank (64 sub-bands, center frequencies 0-20,000 Hz). We then selected 80 sounds, 20 in each of four semantic categories, minimizing within-category repetition (i.e., of multiple sounds from the same objects or animals). To test for categorical structure in low-level features, we first created a matrix comprising all pairwise euclidean distances between stimulus cochleagrams. Categorical membership distances were then computed by contrasting the average between- and within-category matrix subsections. These distances were compared to a null distribution of category membership distances (corrected significance threshold p < 0.05), generated by randomly shuffling rows and columns of the matrix 10,000 times, each time computing the pseudo-category membership distances under the null hypothesis that the category labels are interchangeable. Using this procedure, the final stimulus set was adjusted such that no significant categorical differences were observed in the pattern of pairwise distances. A similar test on a spectrogram-based matrix revealed no significant categorical structure. A 2-d multidimensional scaling (MDS)-based visualization of the sounds’ representational geometry is shown in **Supplemental Figure S1.**

### Experimental design and procedure

Participants were familiarized with the sounds prior to the neuroimaging sessions. Each sound was accompanied simultaneously with a written description, such as ‘a horse neighing,’ ‘a trumpet playing,’ ‘a male shouting angrily,’ and ‘howling wind through a city’ (see **Table S1**). Participants could repeat playing a sound until they were familiar with it. This procedure was completed twice during a monitored setting, and once prior to each experimental session. During the neuroimaging experiment, lighting was dimmed, and participants were instructed to keep their eyes closed at all times. Each sound was presented diotically through earphones (i.e., the same waveform in both channels). In detail, for each MEG and fMRI session, participants completed 16 (MEG) or 12-14 (fMRI) runs, each lasting 330 s. Each sound was presented once in each run in random order, and sounds were randomly interleaved with twenty null (no sound) trials and ten oddball (target detection; monotone double beep) trials. Each sound trial (including oddball trials) consisted of a 500 ms sound presentation followed by 2500 ms of silence preceding the next trial. Optseq2 (https://surfer.nmr.mgh.harvard.edu/optseq/; Greve, 2002) was used to generate, optimize, and jitter the presentation of all trials, including null and oddball-detection trials, and therefore some trials contained extended periods of silence preceding the next sound. During the neuroimaging experiment, participants were instructed to press a button in response to the target (oddball) so that their focus was maintained during the entire sound trial. Null and oddball trials were excluded from the main analysis.

### MEG acquisition

We acquired continuous MEG signals from 306 channels (204 planar gradiometers, 102 magnetometers, Elekta Neuromag TRIUX, Elekta, Stockholm) at a sampling rate of 1 kHz, filtered between 0.03 and 330 Hz. Raw data were preprocessed using spatiotemporal filters (Maxfilter software, Elekta, Stockholm) and then analyzed using Brainstorm (Tadel et al., 2011). MEG trials were extracted with 200 ms baseline and 3 s post-stimulus (i.e., 3,201 ms length), the baseline mean of each channel was removed, and data were temporally smoothed with a low-pass filter of 30 Hz. A total of 16 trials per condition was obtained for each session and participant.

### Multivariate analysis of MEG data

To determine the amount of sound information contained in MEG signals, we employed multivariate analysis using linear support vector machine classifiers (SVM; http://www.csie.ntu.edu.tw/~cjlin/libsvm/; Chang and Lin, 2011). The decoding analysis was conducted independently for each participant and session. For each time point in the trial, MEG data were arranged in the form of 306-dimensional measurement vectors, yielding N pattern vectors per time point and stimulus condition (sound). We used supervised learning, with a leave-one-out cross-validation approach, to train the SVM classifier to pairwise decode any two conditions. For this, we first randomly assigned the trials to N = 4 trial subgroups and sub-averaged the trials within each subgroup. For each time point and pair of conditions, N – 1 pattern vectors comprised the training set and the remaining Nth pattern vectors the testing set, and the performance of the classifier to separate the two conditions was evaluated. The process was repeated 100 times with random reassignment of the data to the subgroups; the overall decoding accuracy of the classifier (chance level 50%) was the mean of those 100 iterations. The decoding accuracy was then assigned to a matrix with rows and columns indexing the conditions classified. The matrix is symmetric across the diagonal, with the diagonal undefined. This procedure yielded one 80 × 80 representational dissimilarity matrix (RDM) of decoding accuracies constituting a summary of representational dissimilarities for each time point. Iterating across all time points in the epoch yielded a total of 3,201 MEG RDMs.

### fMRI acquisition

Magnetic resonance imaging (MRI) was conducted at the Athinoula A. Martinos Imaging Center at the MIT McGovern Institute, using a 3T Siemens Trio scanner (Siemens, Erlangen, Germany) with a 32-channel phased-array head coil. We acquired structural images using a standard T_1_-weighted sequence (176 sagittal slices, FOV = 256 mm^2^, TR = 2530 ms, TE = 2.34 ms, flip angle = 9°). For the experimental task, we conducted 12-14 runs per participant in which 456 volumes were acquired for each run (gradient-echo EPI sequence: TR = 750 ms, TE = 30 ms, flip angle = 54°, FOV read = 210 mm, FOV phase = 100%, ascending acquisition, gap = 10%, resolution = 3 mm isotropic, slices = 44). For the localizer task, two runs were acquired for each participant with 440 volumes per run (gradient-echo EPI sequence: TR = 1000 ms, TE = 30 ms, flip angle = 54°, FOV read = 210 mm, FOV phase = 100%, ascending acquisition, gap = 10%, resolution = 3 mm isotropic, slices = 44).

### Multivariate analysis of fMRI data

fMRI data were processed and analyzed using BrainVoyager QX 2.8 (Brain Innovation, Maastricht, the Netherlands). Data preprocessing included slice acquisition time correction, 3D motion correction, temporal filtering (linear trend removal and high-pass filtering set at 3 cycles/run), and Talairach space transformation (Talairach and Tournoux, 1988). For each participant and session separately, data were realigned and slice-time corrected, and then coregistered to the T_1_ structural scan acquired in the first MRI session. fMRI data was not spatially smoothed. We then modeled the fMRI responses to the 80 sounds with a general linear model (GLM). The onsets and durations of each stimulus presentation (excluding null and oddball trials, which were omitted from further analysis after preprocessing) were entered into the GLM as regressors and convolved with a hemodynamic response function. Movement parameters entered the GLM as nuisance regressors. For each of the 80 conditions, we converted GLM parameter estimates into t-values (z-scored) by contrasting each condition estimate against the implicitly modeled baseline.

Next, we conducted a searchlight analysis to reveal similarity structures in locally constrained fMRI activity patterns. For each voxel v in the brain, we extracted fMRI patterns in its local vicinity (a 4-voxel radius) and calculated condition-wise dissimilarity (1 – Spearman’s R), resulting in an 80 × 80 fMRI representational dissimilarity matrix (fMRI RDM). Repeating this analysis voxelwise across the whole brain yielded 40122 RDMs.

### MEG–fMRI Fusion Searchlight with Representational Similarity Analysis

To relate neuronal temporal dynamics with their spatial loci, we used representational similarity analysis (Cichy et al., 2014, 2016a; Kriegeskorte and Kievit, 2013; Kriegeskorte et al., 2008) to link MEG and fMRI signal patterns. The basic idea is that if two sounds are similarly represented in MEG patterns, they should also be similarly represented in fMRI patterns. Comparing MEG and fMRI RDMs in this way links particular locations in the brain to particular time points, yielding a spatiotemporally resolved map of neural representations.

In representational (dis)similarity space, MEG and fMRI patterns become directly comparable. For each time-specific MEG RDM, we calculated the similarity (Spearman’s R) to each fMRI searchlight’s fMRI RDM, yielding a 3-D map of MEG–fMRI fusion correlations. Repeating this procedure across the brain for each millisecond yielded a MEG–fMRI correspondence “movie” indexing spatiotemporal neural dynamics of cortical auditory responses (see **Movie S1**).

### Regions of Interest

Middle Heschyl’s gyrus (HG; TE1.0), posteromedial HG (TE1.1), anterolateral HG (TE1.2), the planum polare (PP), and the planum temporale (PT) were first identified from anatomical volume masks derived from probabilistic volume maps (Desikan et al., 2006; Morosan et al., 2001; Norman-Haignere et al., 2013). The primary auditory cortex (PAC) was defined as a subdivision of HG including the middle and posteromedial branches (TE1.0 and TE1.1). The temporal voice area (TVA) was identified from probabilistic volume maps of voice-sensitive areas (vocal > nonvocal contrast) along the human superior temporal gyrus (Pernet et al., 2015) and was further restricted to exclude voxels overlapping with PAC, PP, PT, and TE1.2; to reflect this exclusion it is referred to as TVAx here. The left inferior frontal gyrus was identified from spatial coordinates provided by Pernet et al. (2015) and converted to Talairach space using the Yale BioImage Suite Package (Lacadie et al., 2008). For details on the functional localization of these ROIs, see **Supplemental Fig. S3**.

For our visually defined ROIs, data from an independent functional visual localizer was analyzed using a general linear model (GLM), accounting for hemodynamic response lag (Friston et al., 1994). Each participant took part in two runs of the visual localizer task. A 7.1-min functional localizer consisting of photographs of various scenes, faces, common objects, and tile-scrambled images was used to localize the parahippocampal place area (PPA), medial place area (MPA), lateral occipital complex (LOC), fusiform face area (FFA) and an area of early visual cortex (EVC). PPA was defined as a region in the collateral sulcus and parahippocampal gyrus (Epstein and Kanwisher, 1998) whose activation was higher for scenes than for faces and objects (false discovery rate, q < 0.05; this threshold applies to all functional regions localized in individual observers; identified in fifteen participants). MPA (Medial Place Area; see Silson et al., 2016) was a functionally defined region overlapping with retrosplenial cortex-posterior cingulate-medial parietal cortex whose activations were higher for scenes than for faces and objects (identified in 15 participants). In accordance with Grill-Spector et al. (2000), LO, a subdivision of the lateral occipital complex (LOC), was defined as a region in the lateral occipital cortex near the posterior inferotemporal sulcus, with activation higher for objects than scrambled objects (identified in 15 participants). The fusiform face area (FFA) was identified as a region in the extrastriate cortex whose activations were higher for faces than scenes or objects (Kanwisher et al., 1997) (identified in 14 participants). Finally, a control region, early visual cortex (EVC), was identified as an area of peak activity over the posterior occipital cortex (contrast: scrambled images > objects; identified in 15 participants) (Lowe et al., 2016; MacEvoy and Epstein, 2011). Following the identification of these functionally defined regions within participants, probabilistic maps were created using the BrainVoyager QX VOI analysis tool to evaluate the spatial consistency of each region across participants, and converted to volume masks over an averaged brain in Talairach space.

### ROI-based fMRI-MEG fusion

To compute MEG-fMRI fusion time courses for ROIs, we extracted voxel responses within each ROI and computed the condition-specific pairwise Pearson distances to create a single fMRI ROI RDM per subject. Next, we averaged ROI RDMs over subjects and then computed Spearman’s R correlations between each ROI and subject-specific whole-brain MEG RDMs over time to yield fMRI-MEG fusion time courses for each ROI per subject. Final time courses were computed by averaging across subjects.

### Quantification of relative model fits and trends

To quantify the relative strengths of the Cochleagram and Category model fits, and operationalize the trends of those fits over space and time, we first linearized the RDM model fit estimates by applying a Fisher-z transformation to the correlation coefficients. We could thus directly compare the *Δz* difference scores *(Semantic Dominance;* SD) between Category and Cochleagram fits within each ROI in the fMRI data, as well as the whole-brain MEG data (Fischer et al., 2011; Rosenthal and Rosnow, 1991). Performing this analysis within data sets (ROIs or whole-brain MEG) held various factors constant, such as number of voxels, local SNR, etc., that would otherwise confound direct comparisons. The resulting relative difference score was then tested for trends across time (MEG) and space (ROIs). For whole-brain MEG, we compared the SD time course against zero, testing for significance using the methods described above.

For ROI-based analyses, we assigned each ROI a rank reflecting its position in a hypothesized sequence, then computed a Spearman rank correlation between ROIs and their SD Δz-score to assess the direction and significance of the trend toward neural coding dominated by categorical vs. acoustic (cochleagram) information (***α*** = 0.05). The SD trend analysis was conducted with two ROI rank assignments based on hypothesized spatial and temporal criteria. First, to assign ranks based on functional anatomy, we differentiated primary auditory cortex based on convergent anatomical, histological and functional criteria (Morosan et al., 2001; Norman-Haignere and McDermott, 2018; Sweet et al., 2005), then ranked non-primary areas progressing from PAC along and beyond the supratemporal plane. We assigned PAC a rank of 1; TE1.2, PP, and PT (all adjacent to PAC) a rank of 2; TVAx (farther along the posterolateral STG) a rank of 3; LIFG a rank of 4; and FFA, PPA, MPA and LOC a rank of 5. EVC was excluded, as no significant MEG-fMRI fusion time points or RDM model correlations were found in that ROI. To test our temporally based hypothesis, we repeated the analysis, this time assigning ROI ranks guided by MEG-fMRI fusion peak latencies. Finally, to assess how well our anatomical and fusion-guided rank correlations tracked the evolution of Semantic Dominance compared to all possible such correlations, we generated a 20,000-sample bootstrapped distribution of ROI-Semantic Dominance rank correlations, assigning ROIs a randomly drawn rank from 1 to 10 (with replacement) for each sample and comparing the empirical correlations against that distribution.

### Statistical Testing

Nonparametric statistical tests were used to assess significance. To obtain a permutation distribution of maximal cluster size, we randomly shuffled the sign of participant-specific data points (1000 times; cluster definition threshold at p = 0.001; cluster size threshold at p = 0.05), averaged data across participants, and determined 4-D clusters by spatial and temporal contiguity at the cluster-definition threshold. Storing the maximal cluster statistic (size of cluster with each voxel equally weighed) for each permutation sample yielded a distribution of the maximal cluster size under the null hypothesis. We report clusters as significant if they were greater than the 95% threshold constructed from the maximal cluster size distribution (i.e., cluster size threshold at *P* = 0.05).

For statistical assessments of time-series peak and onset latencies, we performed bootstrapping tests. To estimate an empirical distribution over peak and onset latencies of time courses, the subject-specific time series were bootstrapped (1000 times) and the 95% confidence interval was defined on the empirical distribution. For peak-to-peak latency comparisons, we obtained the 1000 bootstrapped samples of two peaks and rejected the null hypothesis if the 95% confidence interval of the peak latency differences did not include zero.

## Acknowledgment

This work was funded by Vannevar Bush Faculty Fellowship program by the ONR to A.O. (N00014-16-1-3116). ST was supported by Training Grant 5T32EY025201-03 and SKERI RERC grant 90RE5024-01-00. The study was conducted at the Athinoula A. Martinos Imaging Center, MIBR, MIT. We thank Dimitrios Pantazis for helpful discussion, and Michele Winter for help with stimulus selection and processing.

## SUPPLEMENTAL INFORMATION

**Supplemental Table S1.** A description of each sound in the stimulus set. Prior to scanning, participants were presented with each sound (played for 500 ms) while reading a written description of the sound. Participants were instructed to listen to sounds until they were familiar with the sound and its interpretation.

| Category | Description | Category | Description |
| --- | --- | --- | --- |
| Animal | A bear roaring | Object | A cash register being opened |
| Animal | A bee buzzing | Object | A cymbal strike |
| Animal | A small cat meowing | Object | A dial tone |
| Animal | A cat purring | Object | A drum roll |
| Animal | A cow mooing | Object | A flute playing |
| Animal | Crickets | Object | Oil in a frying pan |
| Animal | A crow crowing | Object | A hairdryer blowing |
| Animal | A small dog barking | Object | An iphone ringing |
| Animal | A donkey braying | Object | A saxophone playing |
| Animal | A duck quacking | Object | A knife chopping |
| Animal | An elephant trumpeting | Object | A laptop starting sound |
| Animal | A frog croaking in the distance | Object | A phone dialing |
| Animal | A goat bleating | Object | A radio with static sound |
| Animal | A horse neighing | Object | A deck of cards being shuffled |
| Animal | A horse snorting | Object | A toy squeaking |
| Animal | A puppy whining | Object | Fabric being torn apart |
| Animal | A rodent chattering | Object | A toilet flushing |
| Animal | A rooster crowing | Object | A trumpet playing |
| Animal | A sheep baaing | Object | A game console playing |
| Animal | A bird singing | Object | A xylophone playing |
| Voice | A young boy saying "Bell is" | Scene | A band playing music at a show |
| Voice | A female saying "Hannah is" | Scene | An ambulance siren in the street |
| Voice | A female saying "There once was" | Scene | An old car engine starting in the street |
| Voice | A female speaking chinese | Scene | A car horn in highway traffic |
| Voice | A female speaking french | Scene | Bells chiming in a large church |
| Voice | A female speaking german | Scene | Fireworks in the sky |
| Voice | A female speaking hindi | Scene | An helicopter flying through the sky |
| Voice | A female whispering | Scene | Lightning striking a field |
| Voice | A female yelling "No" | Scene | A creaking metal door in a warehouse |
| Voice | A kid speaking a foreign language | Scene | A motorcycle passing-by a road |
| Voice | A kid saying an enthusiastic "Yeah!" | Scene | An orchestra playing music |
| Voice | A male shouting angrily | Scene | Piano recital playing in a concert hall |
| Voice | A male saying "Howdy" | Scene | Firetruck siren |
| Voice | A male laughing | Scene | Pool balls clinking in a bar |
| Voice | A male saying "No" | Scene | Heavy rain in the city |
| Voice | A male saying "Rain or" | Scene | A bell in a school hallway |
| Voice | A male speaking italian | Scene | A warning bell for an approaching train |
| Voice | A male speaking russian | Scene | Water bubbling in a hot spring |
| Voice | A male speaking spanish | Scene | Howling wind through a city |
| Voice | A male saying enthusiastically "Yeah" | Scene | An old house with creaking floors |

**Supplemental Figure S1.**
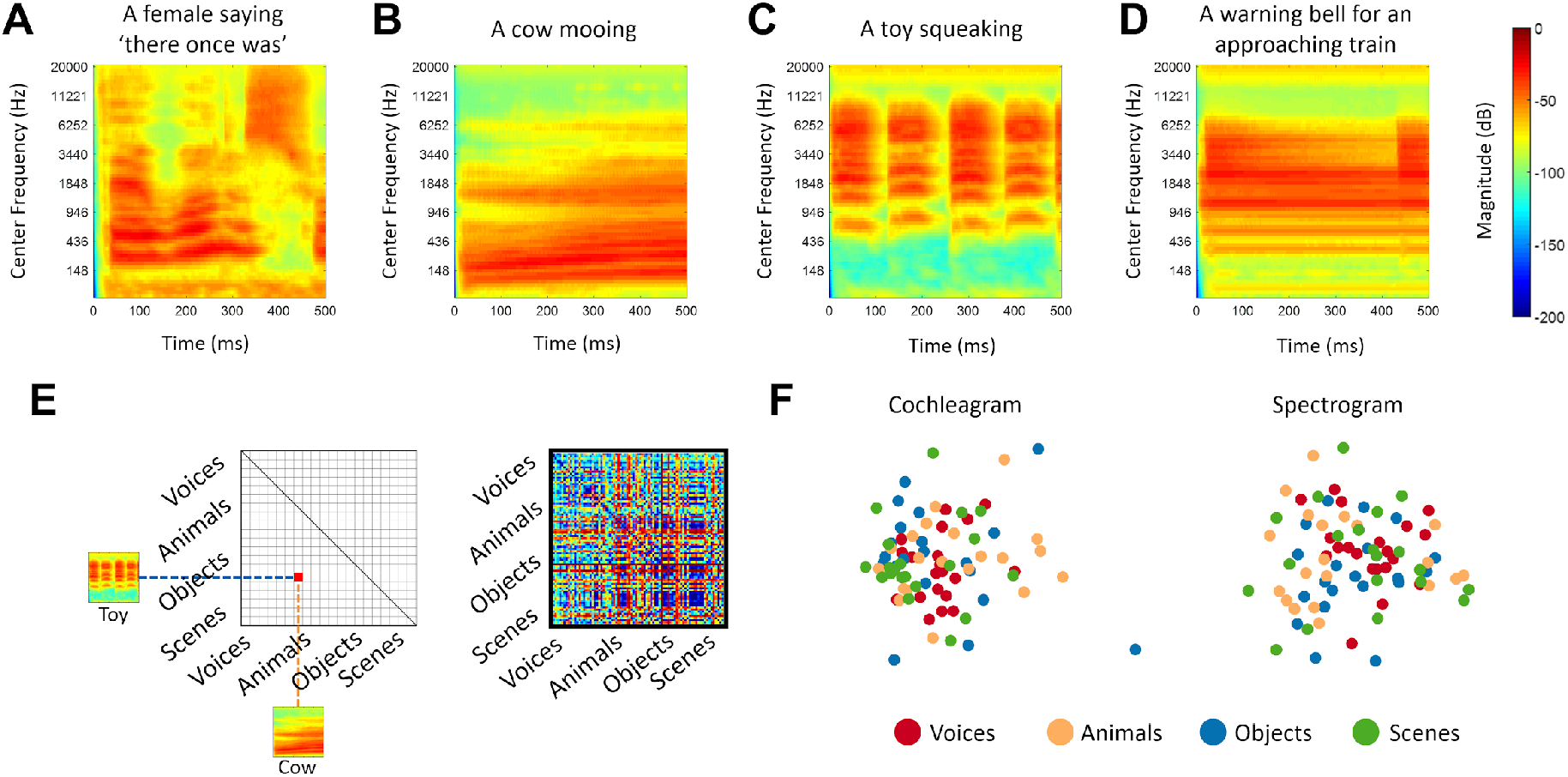
Example stimulus cochleagrams from each category: voice (A), animal (B), object (C), scene (D). Spectral similarity (Euclidean distance) was used to compare all pairs of stimuli, shown schematically in **E** for a sample pair of stimuli *(left)*, and for the entire 80-item stimulus set *(right).* The full matrix was used as the Cochleagram model. **F.** Multidimensional Scaling (MDS) visualization of similarity structures for cochleagrams (*left*) and spectrograms *(right)*, with each dot representing one stimulus exemplar, color-coded by category. To quantify this structure, average between-category distances were compared against between-category distances when category labels were randomly shuffled and permuted 10,000 times. Against a Bonferroni-corrected threshold of p = 0.008, no single category pair differed significantly from randomly permuted category pairs, indicating that categorical structure in brain responses was unlikely to arise from acoustic differences alone.

**Supplemental Figure S2:**
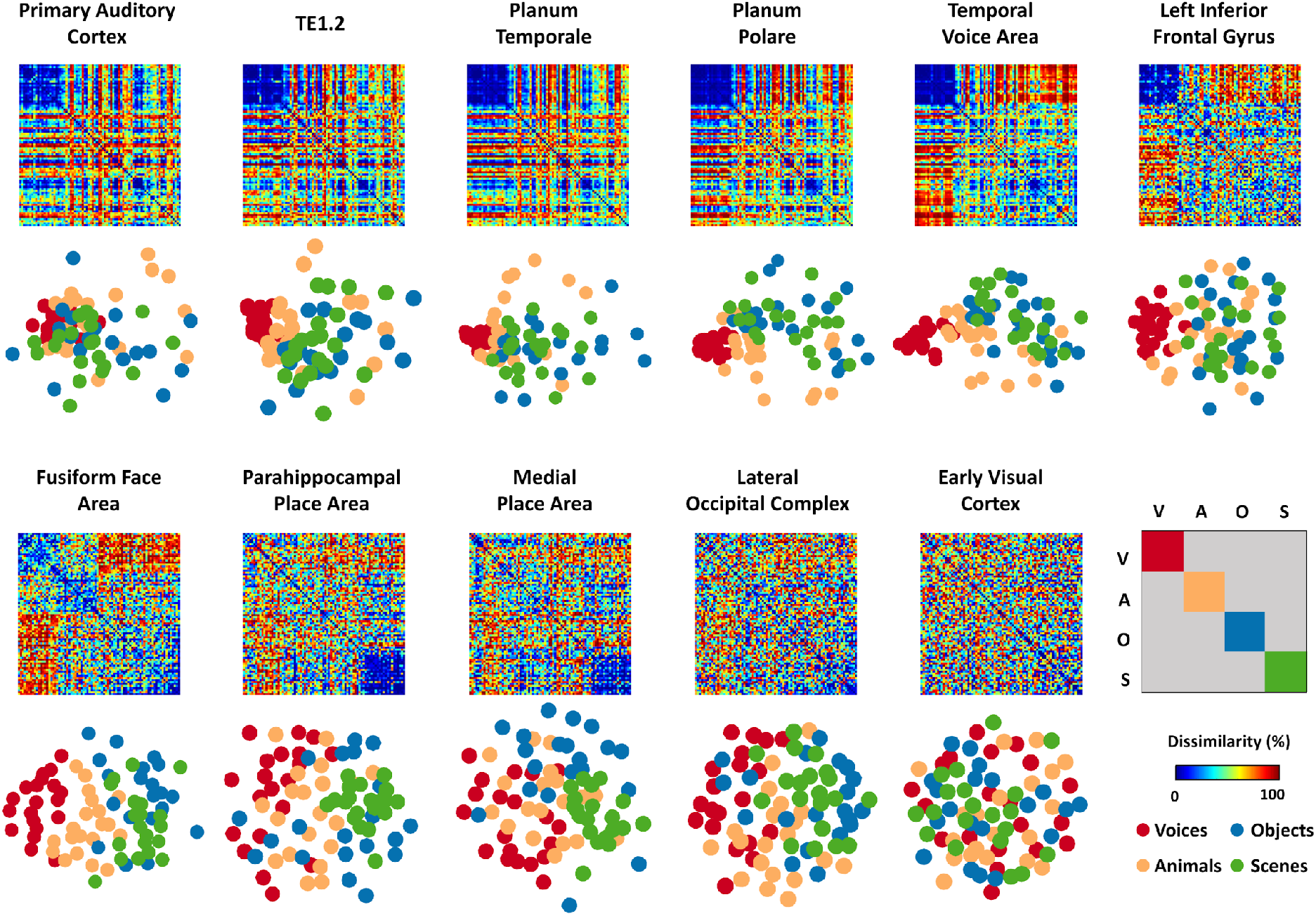
Visualization of Category Selectivity across Regions of Interest. Representational dissimilarity matrices and their corresponding multidimensional scaling visualization (in the first two dimensions) in the 11 fMRI ROIs. Note: Temporal Voice Area refers to TVAx (see main text Methods for details). Each of the 80 stimuli is color-coded by category, as shown in the lower right panel.

### Visualization of category structure using multidimensional scaling

To visualize underlying patterns contained within the complex high-dimensional structure of the 80 x 80 MEG decoding matrix, we used multidimensional scaling (MDS) (Kruskal and Wish, 1978) to plot the data into a two-dimensional space of the first two dimensions of the solution, such that similar conditions were grouped together and dissimilar conditions far apart. MDS is an unsupervised method to visualize the level of similarity contained in a distance matrix (here the decoding matrix; euclidean distance in Fig. S1), whereby conditions are automatically assigned coordinates in space so that distances between conditions are preserved.

**Supplemental Figure S3.**
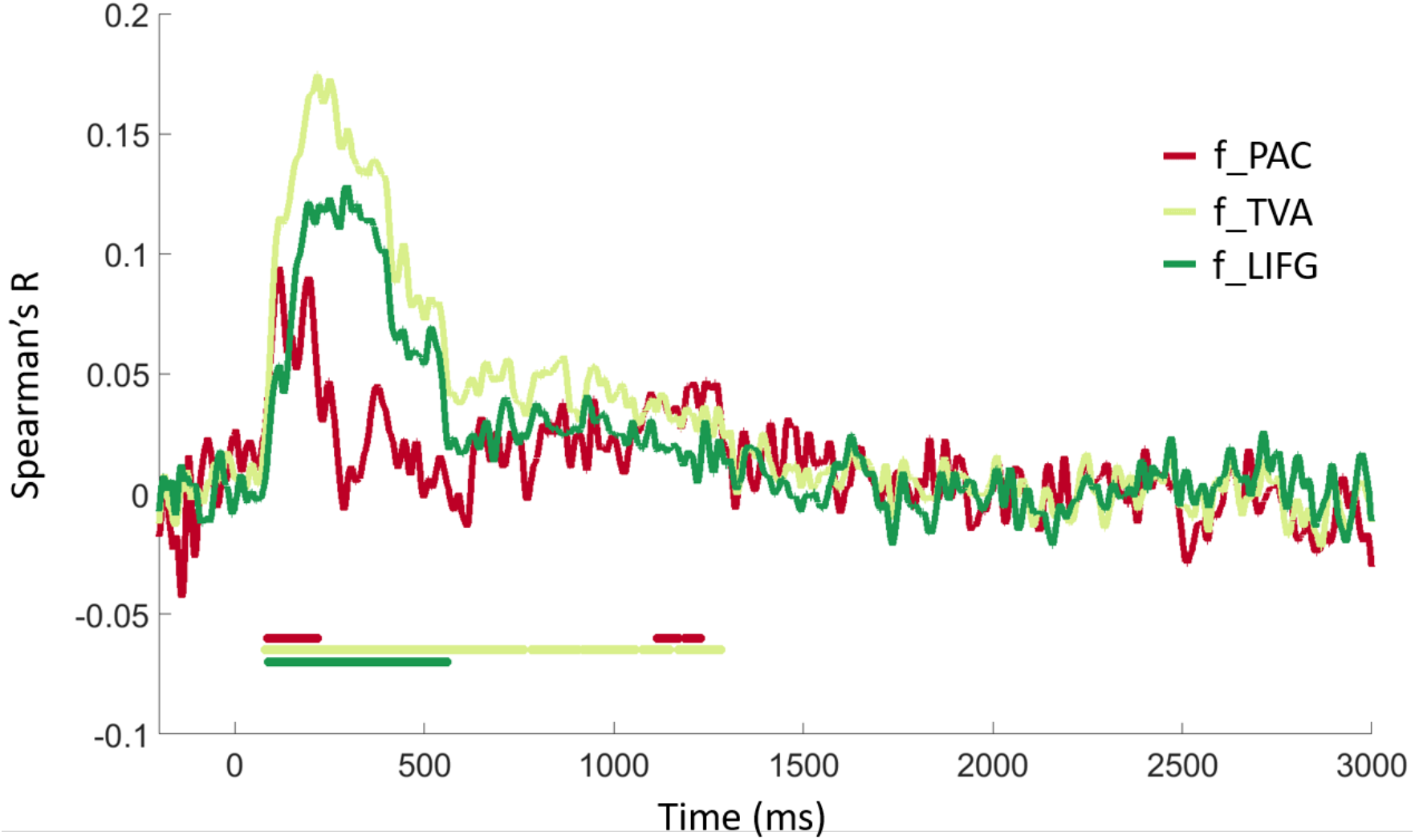
ROI-based fusion of MEG-fMRI in functional auditory ROIs. (using Spearman’s R) in 3 regions of interest, including a functionally defined primary auditory cortex (f_PAC), functionally defined temporal voice area (f_TVA), and left inferior frontal gyrus (f_LIFG). Solid horizontal lines represent significant time points observed, color-coded by region. All statistics, P<0.01, C<0.05, 1000 permutations.

### ROI-based analysis in functionally defined auditory regions

Independent functional data was used to localize three auditory regions of interest. This data was taken from the final run of each fMRI session for each participant, and these runs were not included in the experimental analysis. Primary auditory cortex (PAC) was identified bilaterally in fourteen participants as a region of peak activity (contrast: oddball > null) located over Heschl’s gyrus. Following Pernet and colleagues (2015), the posterior cluster of the temporal voice area (TVA) was identified bilaterally in twelve participants as a region of peak activity (contrast: vocal > non-vocal) located over the middle/posterior superior temporal sulcus. Similarly, a voice area within the extended voice processing network was identified in nine participants as a region of peak activity (contrast: vocal > non-vocal) over the left inferior frontal gyrus (LIFG). Probabilistic mapping was then used to create volume masks over an averaged Talairach brain using BrainVoyager QX.

